# Relationing of Labellum and Glossae Biomorphological characteristics of honeybee workers as breeding tool on honey collection potential of *Apis mellifera* L. honeybee colonies

**DOI:** 10.1101/2021.04.13.439606

**Authors:** Aneela Iqbal, Muhammad Khalid Rafique, Rashid Mahmood, Mamoona Noreen, Ghulam Sarwar, Ishaq Ahmad, Anjum Shahzad, Muhammad Jahangir Shah, Agha Mushtaque Ahmed, Muhammad Akbar Lashari

## Abstract

This study focused on the correlation of honey collection Potential and the length and width of labellum and glossae in worker honey bees (*Apis mellifera* Ligustica). Sixty honeybee *A. mellifera* L. colonies were selected, among these 60 colonies, 3 worker bees were sampled from each colony total numbers of samples collected were 180 adult worker foraging bees. Fifteen colonies for each group were used to check the correlation of honey production with length of labellum, width of labellum, length of glossae and the width of glossae respectively. These worker bees were bought to the laboratory frozen, boiled, dissected and mounted on the slides. Measurements of the labellum length, labellum width, glossae length and glossae width were taken by the stereomicroscope with ocular micrometer at 0.8X magnification. Correlation values for the honey collectionand length and width of labellum and glossae were high and positive. These Results support the perception that worker bees with larger labellum and glossae have more ability for honey collection Potential. It is concluded that Biomorphological characters of labellum and glossae are significantly correlated with the honeycollection Potential in *A. mellifera* L.

## INTRODUCTION

Western honey bees (*Apis mellifera* L.) are social insects, best model for the behavior study and most flourishing specie in Kingdom Animalia [1].Honey bees are very important due their products honey, pollen, bee bread, propolis (vegetable origin), royal jelly bees wax and bee venom (bee secretions) [2]. Honey bees are also very important due to as a trade product, usage in medicine and food and their role in pollination [3].

Honey production is one of the most complex behavioral characters of honey bees that influenced by the morphological strength of honey bees. Study of correlation between the morphological characters and the productivity is of great importance for the future selection of colonies to increase productivity. Study of correlation makes it easier to measure the direction of relationship of morphology and productivity [4].

Morphological characters of honey bees are very important in correlation to honeycollection Potential. Morphological strength of honey bees affects their productivity directly or indirectly. Thereis a positive correlation in the honey production and proboscis size [5].Proboscis of Honeybees consists of following parts; glossae and labellum-honey spoon.Segments elongation of proboscis part’ glossae of honey bees is positively correlated with intake of nectar [6]. For maximum nectar collection honey bees needs long glossae to insert deep into the flower corolla[7,8].

This study has aimed to examine the correlation of honey collection potentialand length and width of labellum and glossae in the western honey bees *Apis mellifera* Ligustica.

## MATERIALS AND METHODS

Sixty beehives undergo queen replacement were screened in the project apiaries for the sample collection at HBRI (Honey bee Research Institute), NARC (National Agricultural Research Centre), Islamabad, Pakistan.

Three worker bees were sampled from each selected colony. Total Sampled bees were 180, brought to the laboratory and kept deep freezer to kill bees. Samples were boiled in water for 2-3 minutes to soften the bees and ethanol was added in bottle of samples. Under magnifying glass lamp proboscis of worker bees were dissected by fore sips and mounted on the double glass slides as show in (figs 2, 4, 6 and 8). After 2-3 h measurements were taken by binocular SZX7 stereo microscope with ocular micrometer at zoom ratio 0.8X ^5^.

**Figure 2:**
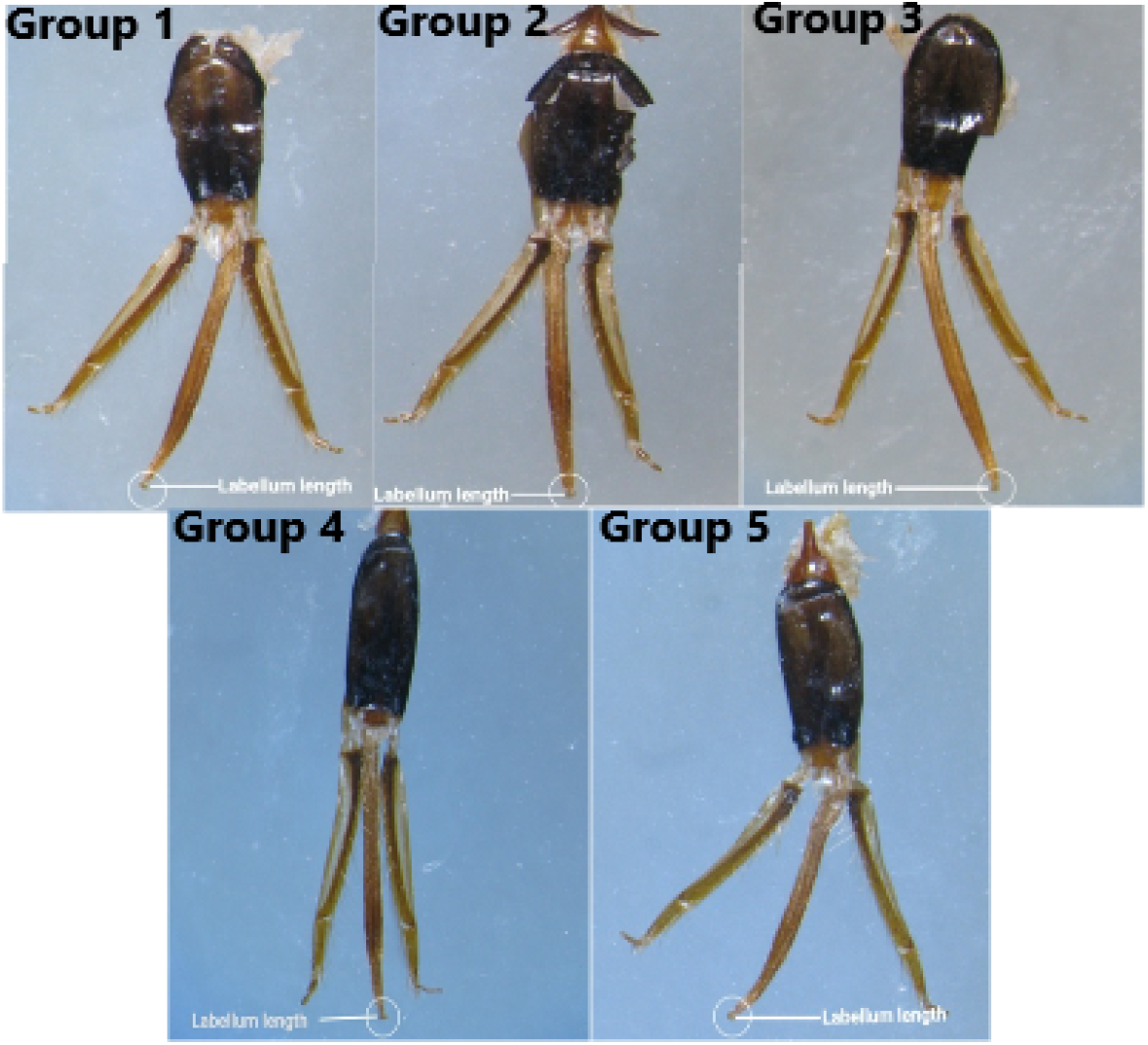
Morphometric Analysis of Proboscis for Labellum Length (Group 1-5)

Colonies were categorized on the base of length and width of labellum and glossae in 5 groups; 3 colonies in each group, after taking the measurements. For spring season colonies were shifted to Margalla hills. Honey was collected and correlation was checked either the correlation is positive or not. Data was analyzed statistically by R-2.6 version statistical software and arranged in Microsoft excel.

**Figure 1:** Steps for the Morphometric analysis of labellum and glossae and their effect on honey collection potential

## RESULTS

### Correlation of labellum length with honey collection Potential

Correlation between the honey collection Potentialand the labellum length are shown in table 1 and (fig 3). Honey yield recorded was 11.08±0.58 kg at the labellum length of 1.59±0.05 and 5.833±0.58 kg at the labellum length of 0.66±0.0. So the results shown that as the length of labellum is high, honey collection increased. This shows that correlation between the honeycollection Potential and labellum length is highly significant P < 0.01.

**Table 1:** Mean of Labellum length in µm (± Standard Error) comparison with honey Collection in kg per hive (± Standard Error) in *Apis mellifera* L.

| Groups | Labellum Length Categories ($\mu\text{m}$) | Labellum Length($\mu\text{m}$) ± S.E | Honey yield (kg) /hive± S.E |
| --- | --- | --- | --- |
| 1 | 0.6 to 0.7 | 0.66±0.0D | 5.833±0.58C |
| 2 | 0.8 to 0.9 | 0.87±0.04C | 7±0.0BC |
| 3 | 1 to 1.05 | 1.02±0.03BC | 8.17±0.58BC |
| 4 | 1.06 to 1.1 | 1.17±0.03B | 9.33±0.58AB |
| 5 | 1.2 to 1.6 | 1.59±0.05A | 11.08±0.58A |
| **F value** |  | 76.3 | 15.3 |
| **P value** |  | 0 | 0.0003 |
| **CV value** |  | 6.57 | 10.91 |

**Figure 3:**
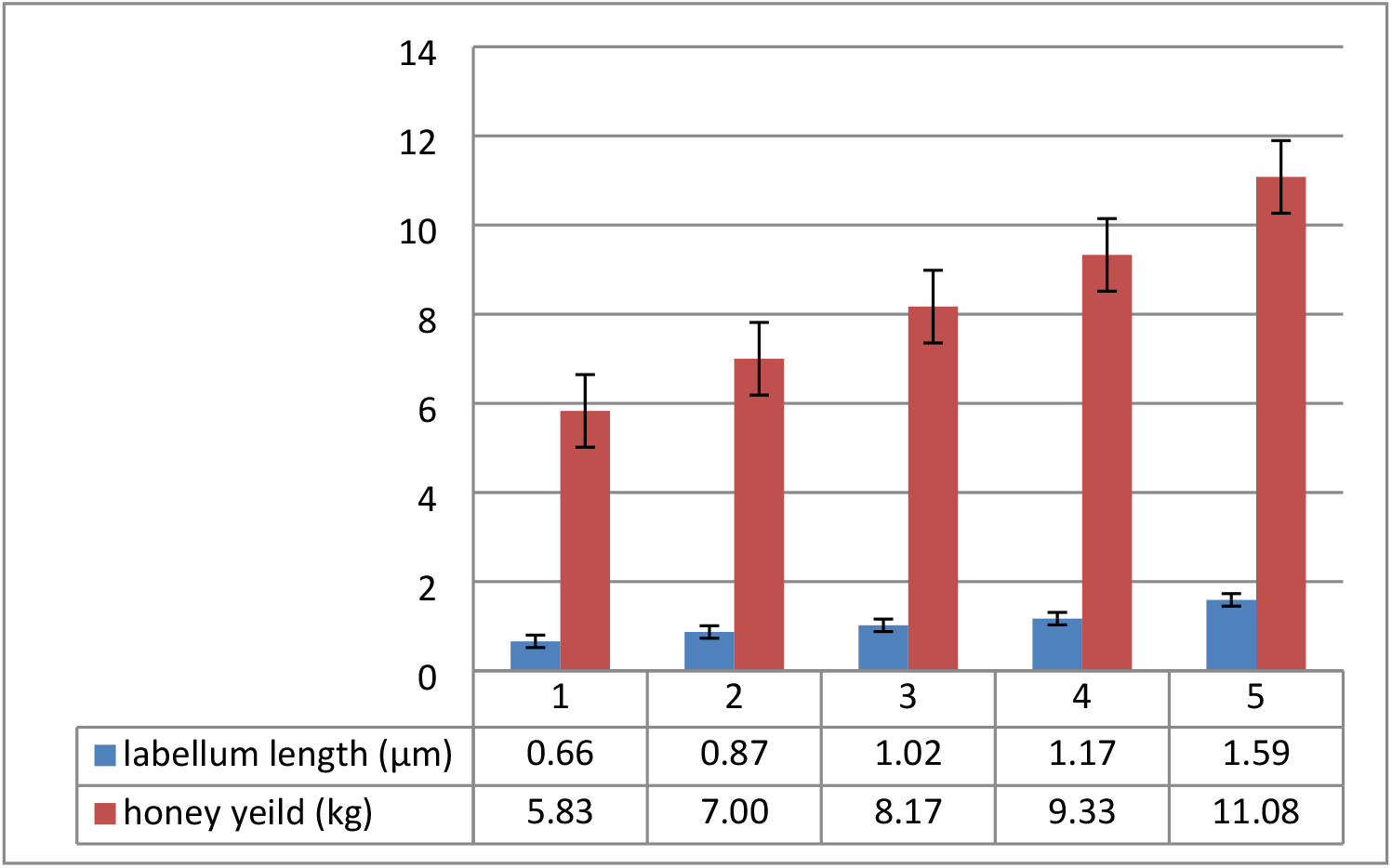
Effect of labellum length variation on honey collection potential.

**Figure 4:**
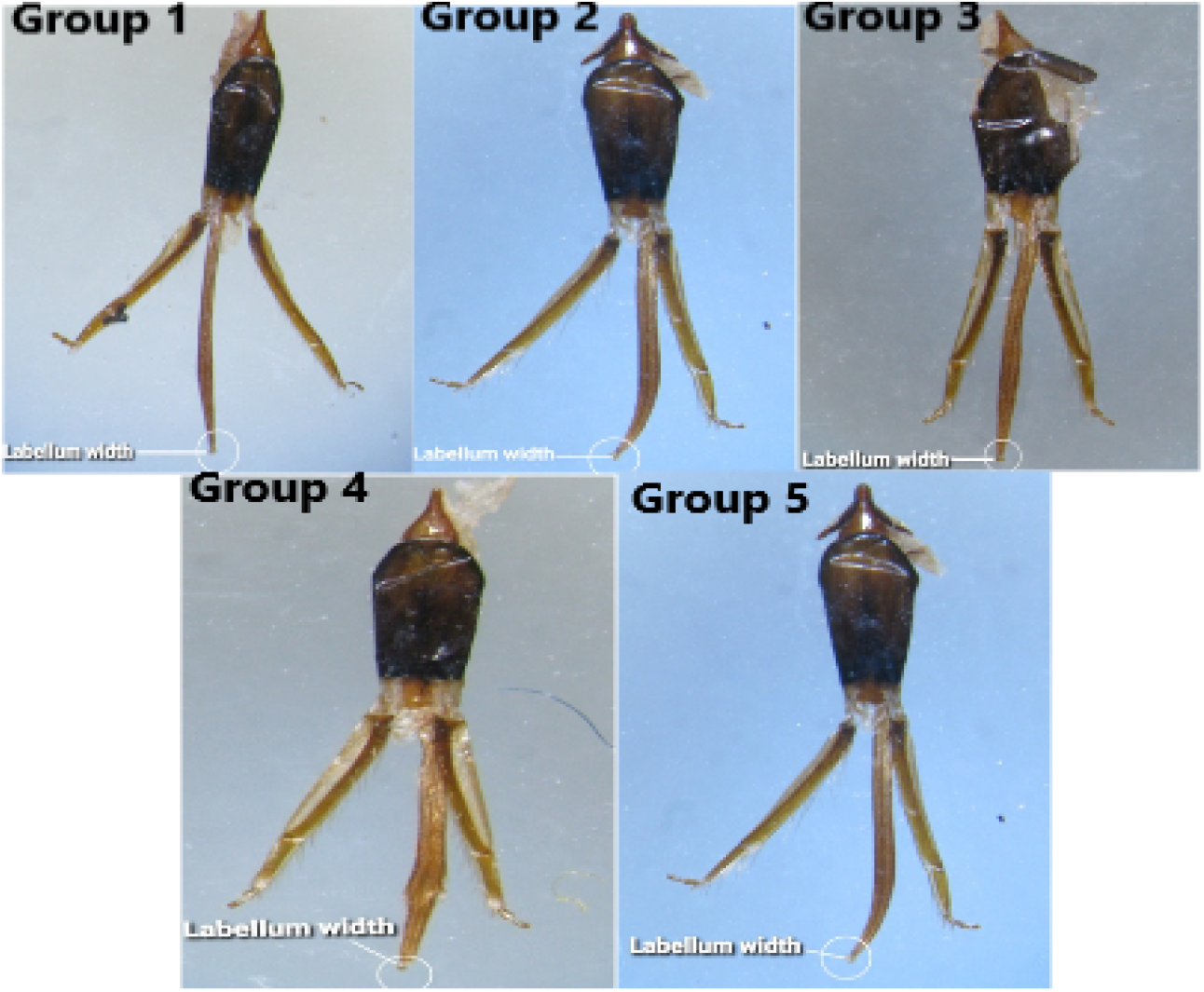
Morphometric Analysis of Proboscis for Labellum Width (Group 1-5)

### Correlation of labellum Width with honey collection Potential

Correlation between the honey collection Potential and the labellum width are shown in table 2 and (fig 3). Honey yield recorded was 11.666±0.58 kg at the labellum width of 1.1553±0.029 and 7±0.00 kg at the labellum width of 0.644±0.06. So the results shown that as the width of labellum is high, honey collectionincreased. This shows that correlation between the honey collection Potentialand labellum width is highly significant P < 0.01.

**Table 2:** Mean of labellum width in µm (± Standard Error) comparison with honey yield in kg per hive (± Standard Error) in Apis mellifera L. (N=3 for each)

| Groups | Labellum Width ($\mu\text{m}$)<br>Categories | Labellum Width<br>($\mu\text{m}$) ± S.E | Honey yield<br>(kg)/hive± S.E |
| --- | --- | --- | --- |
| 1 | 0.55to 0.65 | 0.644±0.06C | 7± 0.0C |
| 2 | 0.75 to 0.85 | 0.833±0 | 8.75± 0.0B |
| 3 | 0.86 to 0.95 | 0.872666±0.03B | 8.75±0B |
| 4 | 0.96 to 0.99 | 0.97866±0.010B | 9.9166±0.58B |
| 5 | 1 to 1.5 | 1.1553±0.029A | 11.666±0.58A |
| **F value** |  | 31.1 | 21.7 |
| **P value** |  | 0 | 0.0001 |
| **CV value** |  | 6.53 | 6.93 |

### Correlation of Glossae length with honey collection Potential

Correlation between the honeycollection Potential and the glossae length are shown in table 3 and (fig 4). Honey yield recorded was 10.5±0.00 kg at the glossae length of 24.0±0.288 and 5.83±0.58 kg at the glossae length of 20.66±0.66. So the results shown that as the length of glossae is high, honey yield increased. This shows that correlation between the honeycollection Potential and glossae length is highly significant P < 0.01.

**Table 3:** Mean of Glossae length in µm (± Standard Error) comparison with honey yield in kg per hive (± Standard Error) in Apis mellifera L. (N=3 for each)

| Groups | Glossae length ($\mu\text{m}$)<br>Categories | Glossae length<br>($\mu\text{m}$) ± S.E | Honey yield<br>(kg)/ hive ± S.E |
| --- | --- | --- | --- |
| 1 | 19.3 to 21.2 | 20.66±0.66C | 5.83±0.58C |
| 2 | 21.3 to 22.6 | 22.22±0.22BC | 7.583±0.58B |
| 3 | 22.7 to 23.0 | 22.9±0.1AB | 8.75±0.0B |
| 4 | 23.1 to 23.2 | 23.142±0.021AB | 8.75±0.0B |
| 5 | 23.3 to 24.0 | 24.0±0.288A | 10.5±0.0A |
| **F value** |  | 13.3 | 21.8 |
| **P value** |  | 0.0005 | 0.0001 |
| **CV**<br>**value** |  | 2.63 | 7.71 |

### Correlation of Glossae Width with honey collection Potential

Correlation between the honey production and the glossae width are shown in table 4 and (fig 5). Honey yield recorded was 11.666±0.58 kg at the glossae width of 1.94±0.053 and 5.83±0.58 kg at the glossae width of 1.33±0.05. So the results shown that as the width of glossae is high, honey yield increased. This shows that correlation between the honey collection Potentialand glossae width is highly significant P < 0.01.

**Table 4:**
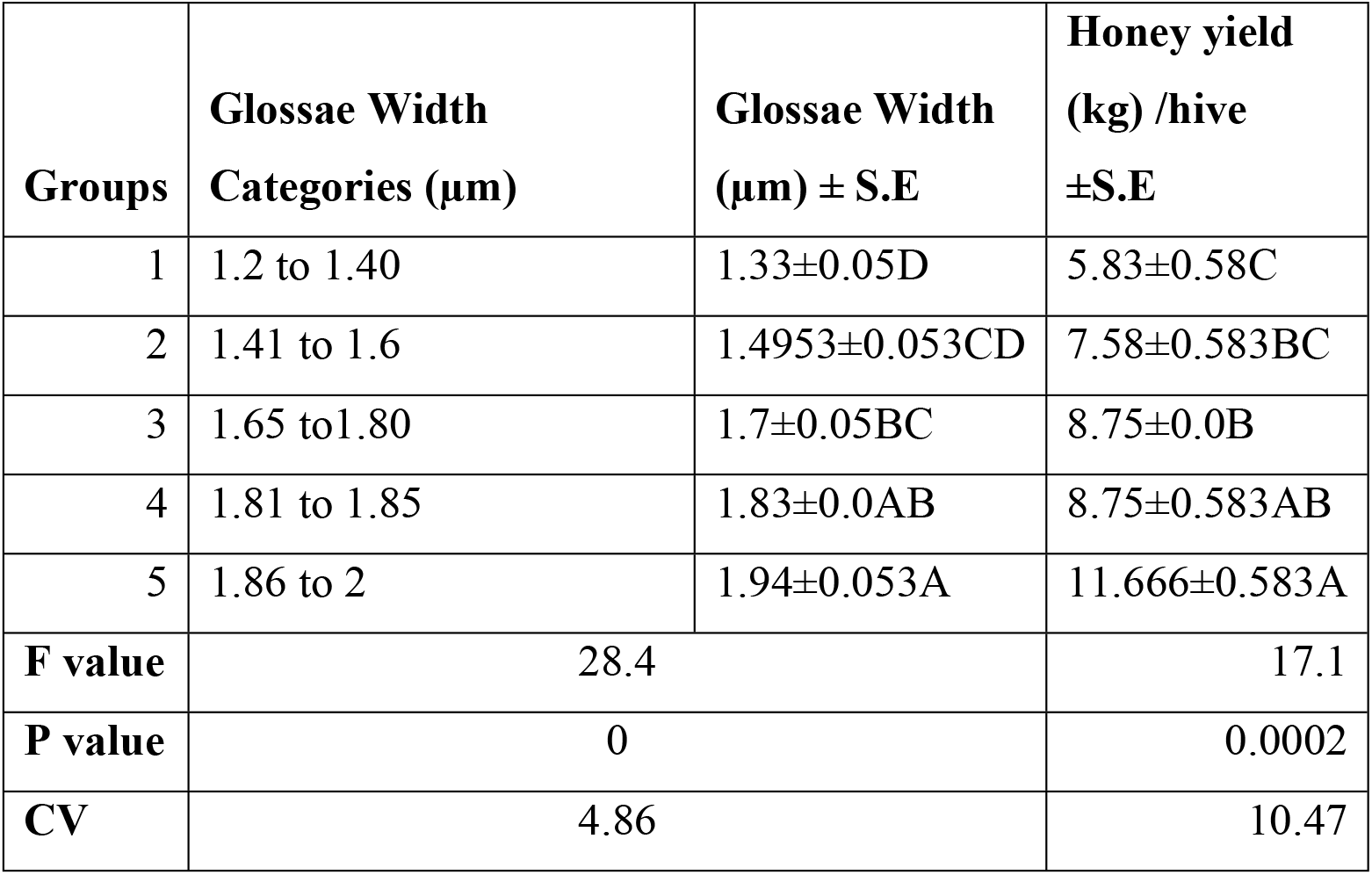
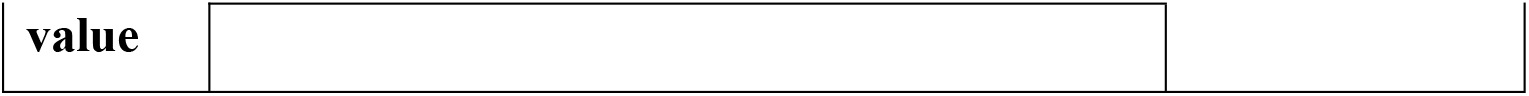
Mean of Glossae width in µm (± Standard Error) comparison with honey yield in kg per hive (± Standard Error) in *Apis mellifera* L. (N=3 for each)

| Groups | Glossae Width Categories ($\mu\text{m}$) | Glossae Width ($\mu\text{m}$) ± S.E | Honey yield (kg) /hive ±S.E |
| --- | --- | --- | --- |
| 1 | 1.2 to 1.40 | 1.33±0.05D | 5.83±0.58C |
| 2 | 1.41 to 1.6 | 1.4953±0.053CD | 7.58±0.583BC |
| 3 | 1.65 to1.80 | 1.7±0.05BC | 8.75±0.0B |
| 4 | 1.81 to 1.85 | 1.83±0.0AB | 8.75±0.583AB |
| 5 | 1.86 to 2 | 1.94±0.053A | 11.666±0.583A |
| **F value** | 28.4 |  | 17.1 |
| **P value** | 0 |  | 0.0002 |
| **CV** | 4.86 |  | 10.47 |

**Figure 5:**
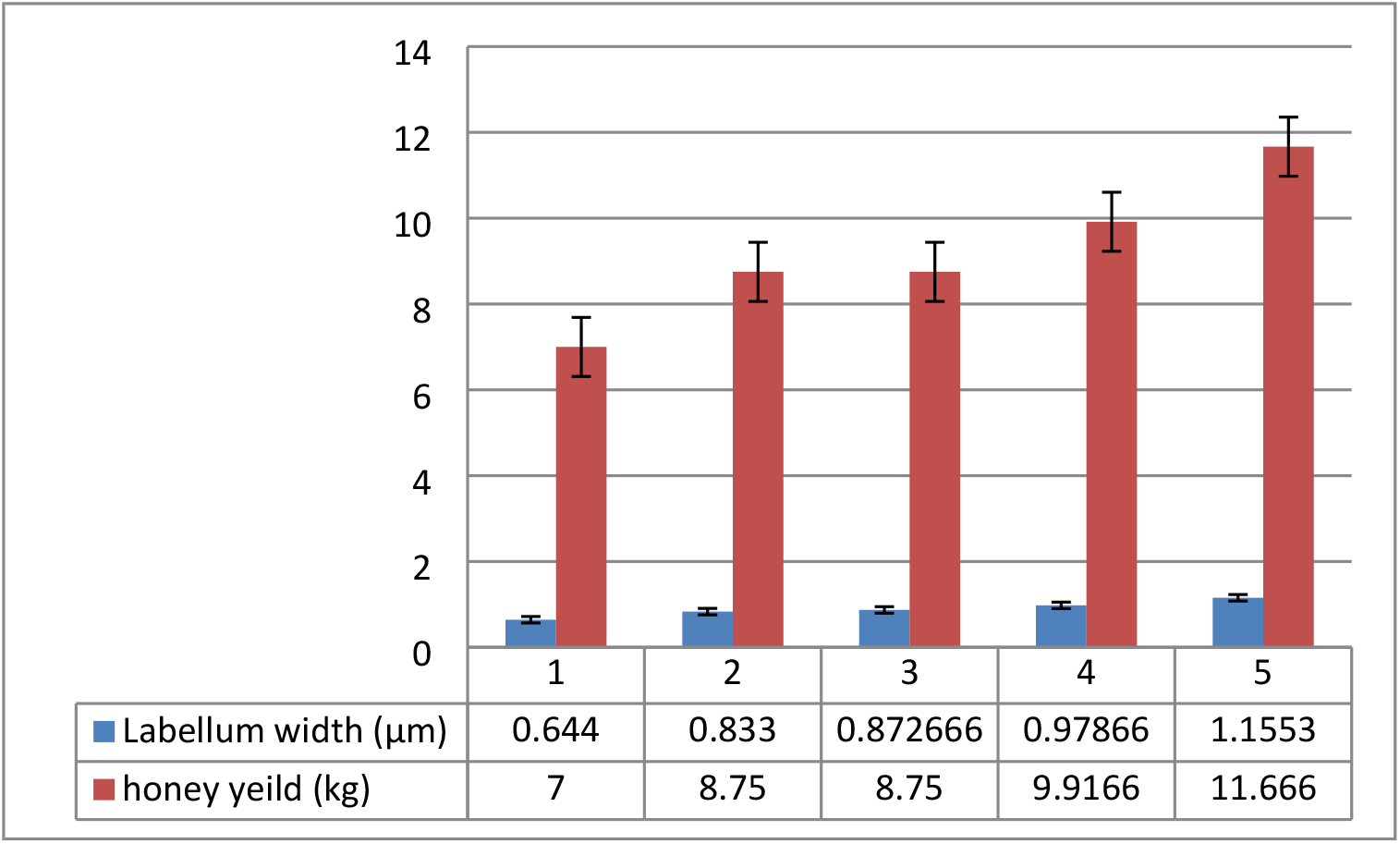
Effect of labellum width variation on honey collection potential.

**Figure 6:**
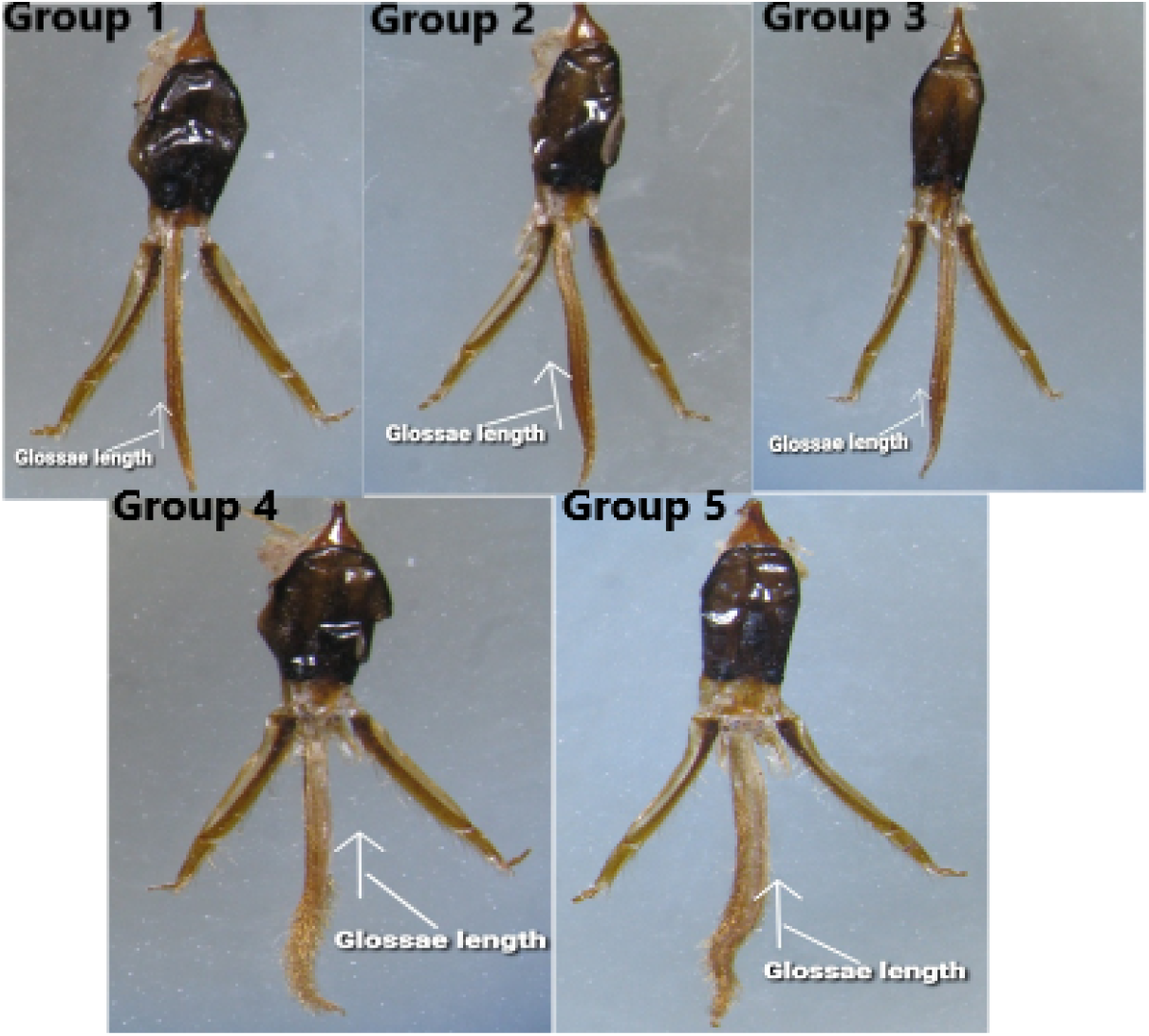
Morphometric Analysis of Proboscis for Glossae length (Group 1-5)

**Figure 7:**
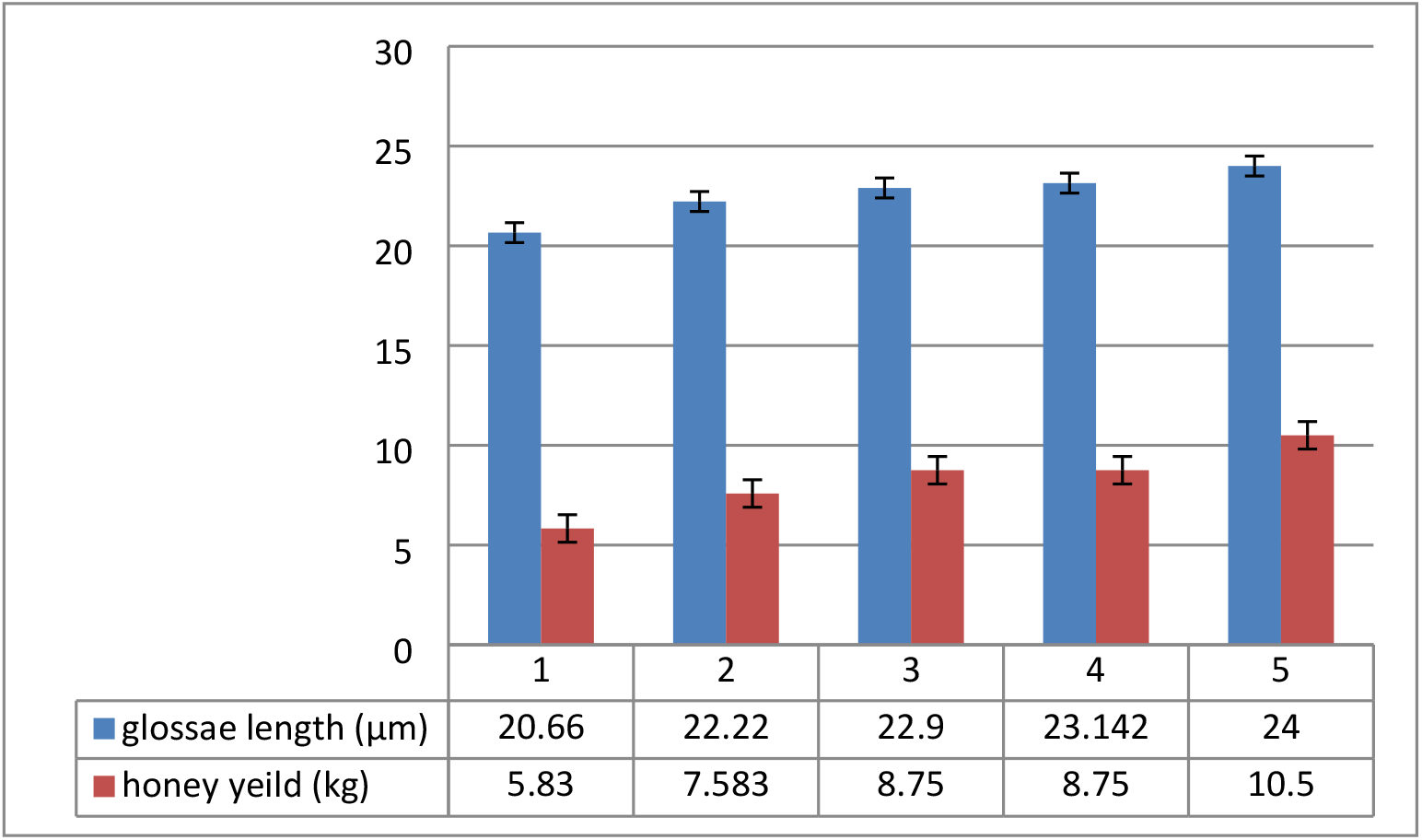
Effect of glossae length variation on honey collection potential.

**Figure 8:**
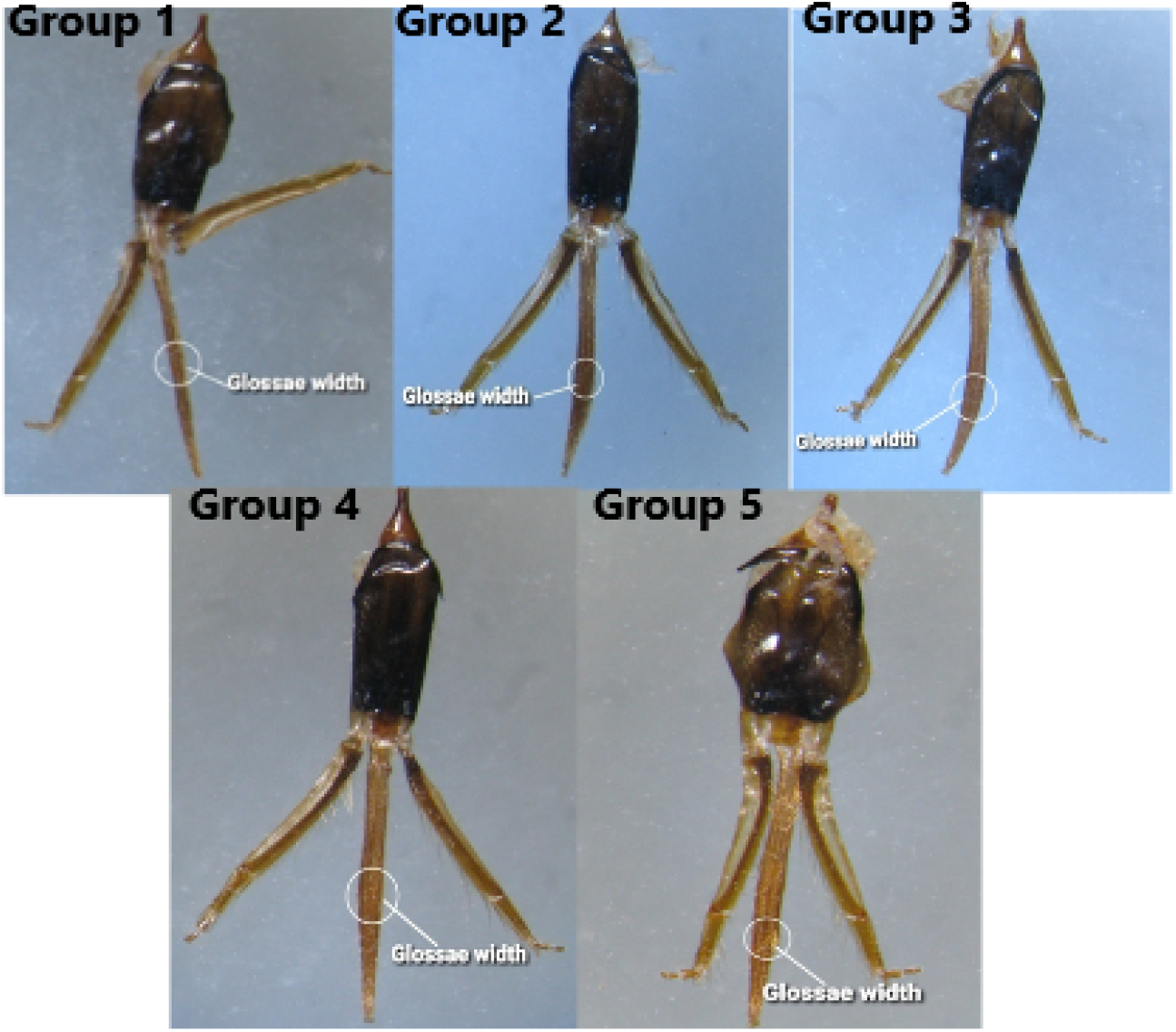
Morphometric Analysis of Proboscis for GlossaeWidth (Group 1-5)

**Figure 9:**
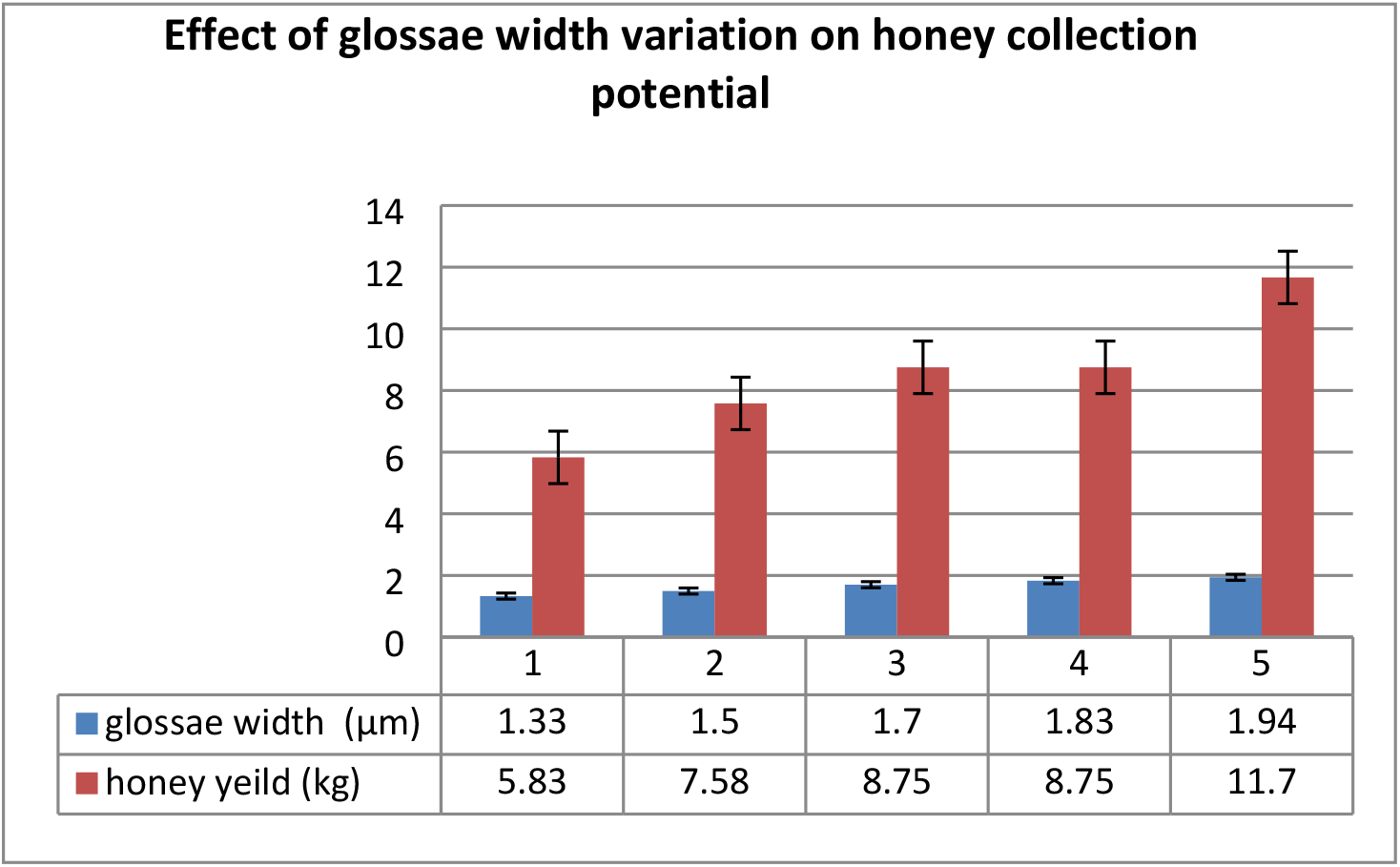
Effect of glossae width variation on honey collection potential.

## DISCUSSION

Globally including Pakistan honey became an important product for trade due to its rising preference as a natural food, its usage in medicine and as medicine [9].

Preceding studies regarding honey production have confirmed that the morphology of proboscis of honey bees are positively correlated to honey collection potential[10]. In Bombus it was observed that length of proboscis is correlated with the corolla’s depth of flowers [11].Honey production is correlated to overall size of honeybees [12].

Morphological strength of wings and legs also influenced the collection potentialof honey. As strong the wings and legs are honeybees could gather nectar and pollen far from the colony [13].

So for the prediction of healthy and productive colony morphological strength of body including tibia length, metatarsus width, Proboscis length, femur length, length and width of fore wing, hind wing length could be important parameters [5,10].

This experimental study was conducted on *Apis mellifera* L. for the prediction of correlation of honey collection potentialand labellum length, honey collection potentialand labellum width, honey collection potentialand glossae length and honey collection potentialand glossae width. Results showed that the correlation between the honey collection potentialand labellum length, labellum width, glossae length and glossae width is highly significant P < 0.01.

These results show the significant F-values for the after analysis of variance for the labellum length, labellum width, glossae length and glossae width in correlation to honey collection potentialas in the earlier study of [14]. F-Value for labellum length was 76.3 and F-value for honey yield correlation with the labellum length was 15.3. F-Value for labellum width was 31.1 and honey yield correlation with the labellum width was 21.7. F-Value for glossae length was 13.3 and F-Value for honey yield correlation with the glossae length was 21.8. F-Value for glossae width was 28.4 and F-Value for honey yield correlation with the glossae width was 17.1.

## CONCLUSION

It is concluded from the data obtained and analyzed that Morphological characters of labellum and glossae are significantly correlated with the honey collection potentialin *A. mellifera* L. So for the selection of best breeder colony, length and width of labellum and glossae could be the best parameters like other traits of the colony.

## ACKNOWLEDGEMENT

This research work was supported by ALP – project entitled “Superior Quality honeybee queen production through Non-traditional Techniques” under Grant No. ALP NR-047 of Honeybee Research Institute (HBRI), National Agricultural Research Centre (NARC), Pakistan Agricultural Research Council (PARC), federal Ministry of National Food Security and Research Islamabad (MNFS & R) Pakiostan.

## REFERENCES

1 Hinojosa, I. & Rasnitsyn, A. P. A honey bee from the Miocene of Nevada and the biogeography of Apis (Hymenoptera: Apidae; Apini). Proc Calif Acad Sci. (2009).

2 Bargańska, Ż., Ślebioda, M. & Namieśnik, J. Honey bees and their products: Bioindicators of environmental contamination. Critical Reviews in Environmental Science and Technology 46, 235–248 (2016).

3 Ediriweera, E. & Premarathna, N. Medicinal and cosmetic uses of Bee’s Honey–A review. Ayu 33, 178 (2012).

4 Souza D. C., Cruz C. D., Campos L. A. d. O. & Regazzi A. J. Correlation between honey production and some morphological traits in africanized honey bees (Apis melifera). Ciência Rural 32, 869–872 (2002).

5 Abou-Shaara, H. F., Al-Ghamdi, A. A. & Mohamed, A. A. Body morphological characteristics of honey bees. Agricultura 10, 45–49 (2013).

6 Zhu, R. et al. Feeding kinematics and nectar intake of the honey bee tongue. Journal of Insect Behavior 29, 325–339 (2016).

7 Krenn, H. W., Plant, J. D. & Szucsich, N. U. Mouthparts of flower-visiting insects. Arthropod Structure & Development 34, 1–40 (2005).

8 Wilhelmi, A. P. & Krenn, H. W. Elongated mouthparts of nectar-feeding Meloidae (Coleoptera). Zoomorphology 131, 325–337 (2012).

9 Khan, H. U., Anjum, S. I., Sultana, N. & Khattak, B. Honey production potential of the honey bee (Apis mellifera) in Karak and Kohat, Pakistan. (2016).

10 Mostajeran, M., Edriss, M. & Basiri, M. in Seventh World Congress on Genetics Applied to Livestock Production. August. 19–23.

11 Inouye, D. W. The effect of proboscis and corolla tube lengths on patterns and rates of flower visitation by bumblebees. Oecologia 45, 197–201 (1980).

12 Kolmes, S. A. & Sam, Y. Relationships between sizes of morphological features in worker honey bees (Apis mellifera). Journal of the New York Entomological Society, 684–690 (1991).

13 Mostajeran, M., Edriss, M. A. & Basiri M. R. Analysis of colony and morphological characters in honey bees (Apis mellifera meda). Pak. J. Biol. Sci 9, 2685–2688 (2006).

14 Pignata, M. I. B., Stort, A. C. & Malaspina, O. Study of the length of the mouthparts of Africanized, Caucasian and Africanized/Caucasian honey bee crosses, and relationships between glossa size and food gathering behavior. Genetics and molecular biology 21 (1998).

